# Extending digital biology: bacterial survival and morphological heterogeneity under antibiotic stress

**DOI:** 10.1101/2025.11.25.690353

**Authors:** Erik Maikranz, Andrey Aristov, Lena Le Quellec, Charles N. Baroud

## Abstract

The morphology of bacteria is modified by antibiotic stress while also serving to survive the antibiotic. However associating morphological descriptors with quantitative measurements of cell survival remains elusive. Here we present a workflow to generate morphological signatures for the progeny of individual cells for 168 different antibiotic conditions. The workflow uses stationary microfluidic droplets, to encapsulate and grow bacteria, and confocal microscopy to image the contents of each droplet. A custom image analysis pipeline is developed to interact with the images in order to label of the morphologies within a subset of the images and train a neural network. The network yields a multidimensional morphological signature for 82000 droplets, showing the co-existence of different morphologies even for the progeny of individual cells. The morphological signatures are different for varying antibiotic type and concentration, thus providing a way to distinguish antibiotics by their mode of action. By combining these morphological signatures with the digital detection of survival within droplets, this workflow can serve to understand the emergence of antibiotic resistance or to identify antimicrobial activity of unknown substances.

---

Droplet microfluidics has been the enabling technology for a range of biological applications that require the ability to distinguish a variety of single-cell behaviors or that can detect rare events. These high-sensitivity and low detection threshold rely on the ability to produce and measure the contents of a large number of droplets, which provides a precise quantification of the biological assay (*1*). One of the main successes of the droplet technology has been in the development of digital biology, in which the outcome of a biological assay is treated as a binary signal, 0 to indicate a negative result or 1 to indicate a positive result (*2*). This digital approach has been most successful when applied to nucleic acid detection, where droplet based techniques formed the enabling technology for digital PCR (*3–5*).

Beyond PCR, the digital approach has been applied to microbiology and antibiotic sensitivity testing, where digital measurements have provided a way to test the single-cell susceptibility to a given dose of antibiotics (*6–9*). In spite of its statistical power however, this digital approach has only been applied to simple fluorescent signals that can be easily thresholded. It therefore fails to provide any insights to identify which cells are able to survive the antibiotic and how they do so. Obtaining such complex insights would require more information about the cells that do survive.

Among the many metrics used to measure the stress response of microorganisms, morphology has frequently been a reliable marker. This is true on the scale of bacterial colonies, whose morphologies have long been used to identify the species, and is now performed using artificial intelligence (AI) (*10–12*). The morphological signature has also been used on the scale of individual cells, where many approaches have been developed to measure bacterial morphology and relate it to antibiotic action, ranging from the use of electron microscopy (*13–16*) to high-resolution fluorescence images of membranes and nucleoli (*17–20*). The optical microscopy is then combined with image analysis to perform so-called cytological profiling and extract morphological descriptors such as area, length and circularity of individual cells. The motivation in this case is to relate an appropriate subset of these morphological features to the mechanism of action of a variety of antibiotics and to also identify susceptible subpopulation of bacteria (*19, 21, 22*). In parallel mechanistic studies have shown how specific morphologies indicate biological modifications that mediate the ability of cells to escape from antibiotics (*23–25*).

In the current manuscript we extend the digital framework to include the morphology of viable cells under antibiotic stress. For this we grow E. coli cells overnight with antibiotics and observe their shape at high resolution (*9*). An automated workflow and analysis pipeline is developed to obtain a data set containing 82,000 droplets and 168 antibiotic conditions (type and concentration), which are then analyzed using a neural network to label the presence (1) or absence (0) of each morphology in each droplet. The resulting classification yields a multi-dimensional signature of the morphology for the progeny of cells that survive the antibiotic concentration. Below we begin by explaining the experimental and computational pipelines, followed by a description and discussion of the impact of the biological results.

## Results

### Experimental protocol and description of the droplet contents

The microfluidic experiments used in this study follow the protocol of Ref. (*9*). In brief, the microfluidic device consists of a wide (4 mm) and thin (15 *μ*m) chamber, whose ceiling is patterned with 500 microfluidic anchors (see Fig. 1a) (*26, 27*). The chip is first filled with a fluorinated oil, then with a suspension of bacteria and antibiotic. Afterwards the oil is re-introduced into the chip, which leads to a single droplet being formed inside each of the anchors, thus creating 500 individual micro-compartments with a volume of approximately 2 nl each (*27, 28*). The loading of the sample and droplet breaking can be done in under 10 minutes per chip and six chips are loaded in series for every experimental run. Antibiotics are added to the bacterial suspension just before being loaded into the chip so that each of the six chips corresponds to a given antibiotic type and concentration. The chips are then placed on a 3D-printed holder (see Fig. 1b), such that each experimental run tests six antibiotic concentrations with the same starting bacterial culture and conditions.

**Figure 1:**
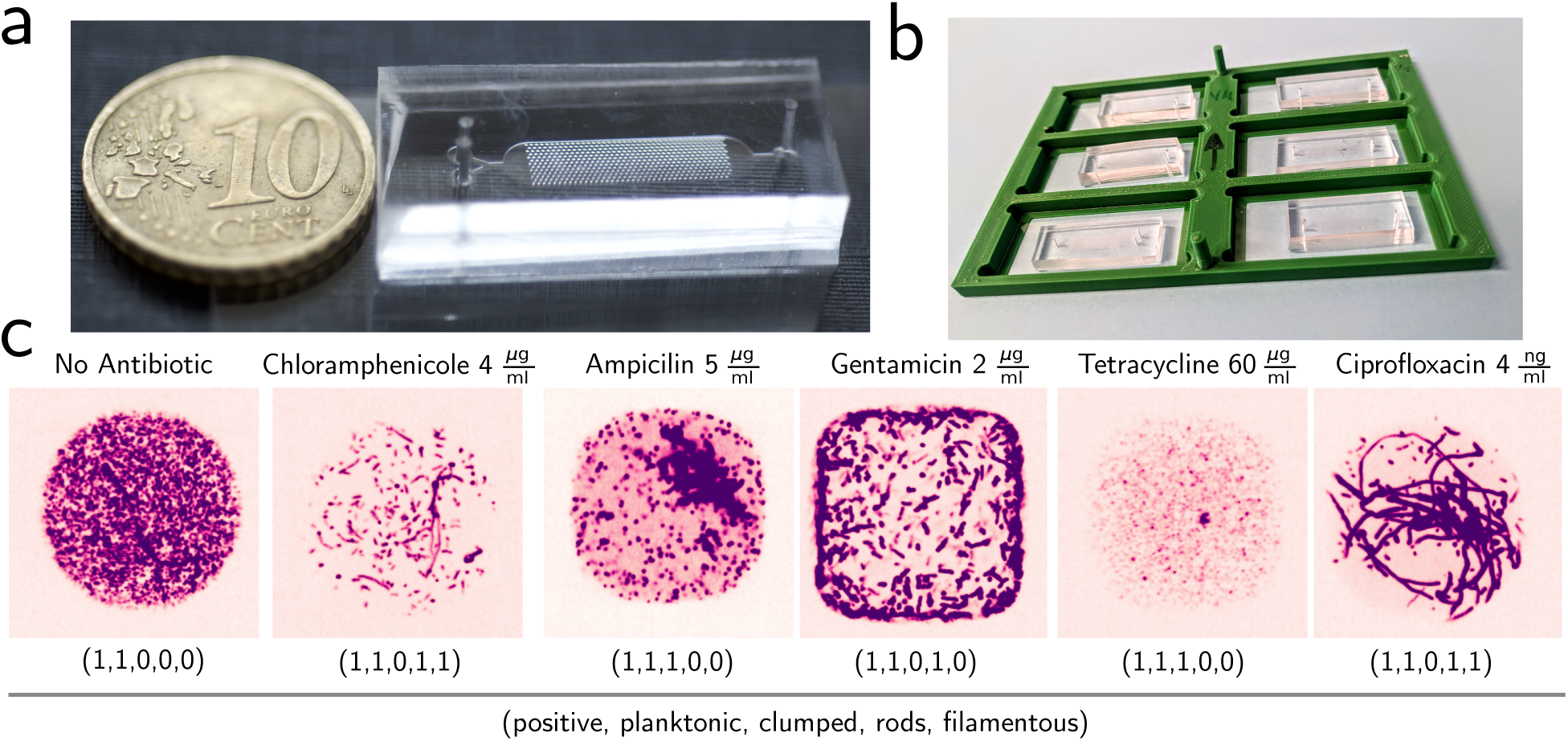
Experimental setup and bacterial morphologies. **(a)** A single microfluidic chip with 500 anchors in the central region. The region of the chip that contains the anchors is 15×4 mm in size and each anchor is 10015×10015×100 *μ*m. The geometry and experiments are detailed in (*9*). **(b)** The individual microfluidic devices are grouped together in a 3D printed chip holder that allows six conditions to be imaged in parallel on the same microscope. **(c)** Different antibiotics lead to different morphologies in the droplets. This figure shows an example droplet for each antibiotic, always at sub-lethal (sub-MIC) concentrations. Five classes are defined: positive, planktonic, clumped, rod-shaped and filamentous cells. The numbers between parentheses correspond to the binary value for the five different classes. Note that these classes are non exclusive for each droplet.

As a bacterial model we use E. coli W3110, labeled with 405 red fluorescent protein (lacYZ:mRFP-1) (*29*), which allows us to image the cells in fluorescence. The chips are imaged a first time just after loading, after which they are placed overnight in an incubator at 37 ^◦^C. A second scan of the chips is then performed on the second day. The imaging on days 1 and 2 begins by acquiring a full scan of each chip in bright-field, followed by a scan in confocal mode through stacks of 25 vertical positions, all with a 20x 0.7 NA objective. The current study included 5 antibiotics (and a non-antibiotic condition), which were tested under multiple concentrations and with several replicates. The resulting data set consisted of 168 microfluidic chips, each of which contained 500 independent droplets on both days, for a total of 168,000 droplet images.

The resolution and sensitivity of the spinning-disk images is sufficient to detect individual bacterial cells within each droplet. A few representative images from maximum projections along vertical positions are shown in Fig. 1c for five different antibiotic conditions, in addition to a no-antibiotic control. Following visual inspection of the droplet contents in many experiments, we introduce five labels to describe these colonies. Foremost, we introduce *”positive”* to denote successful colony formation. The positivity of a droplet was determined by visual inspection, in order to avoid artifacts associated with automatically counting cells that had widely different morphologies. Morphological descriptors then begin with *”planktonic”*, which denotes bacteria that form a suspension of individual cells, each of them having the appearance of a small dot. In contrast, sometimes cells adhere together and form a solid-like mass within the droplet, which we refer to as *”clumped”*. Furthermore, in some conditions we observe *”rod”*-like cells. These cells are longer than the normal planktonic state but do not fill the width of the droplet. Finally, we also identify a *”filament”* morphology, in which cell are long, bent and often fill the entire droplet.

As shown in the representative examples of Fig. 1c, one droplet can often contain a mixture of several morphologies at the same time. For this reason we use non exclusive labelling on the droplet-level. Moreover, the morphologies are not specific to particular antibiotics, as cells cultured in the absence of antibiotics can yield both planktonic or clumps, or a combination of morphologies in different droplets. Similarly, cells cultured with antibiotics can form a variety of colony morphologies in the different droplets within the same chip. Finally, there is no obvious relation between the morphology of the bacteria and their number in a given droplet.

### Data management and dynamic interactions with the images

The images shown in Fig. 1c are representative of a large number of individual droplets under identical microscopy settings. As such they are very well suited for analysis using machine learning tools. However a strong bottleneck arises due to the scale of the data as each experimental run generates 62 GB of data that must be stored on a remote server. As a result the sheer size of these data makes it impossible to interact with them dynamically without significant pre-processing steps.

A pre-processing pipeline is therefore implemented. It begins by reducing the fluorescence stacks from original Nikon nd2 files to max-projected images along the z-axis, which are then concatenated with the 2D brightfield images of the chip and saved into multiscale zarr arrays. Then the droplet anchors are automatically detected using a template matching approach, since their relative positions are known from the microfluidic design (*9*). The detected coordinates of each droplet are saved in a csv file and used to produce cropped images of size 30015×300 pixels for each droplet. These crops are saved into a separate zarr array in full resolution. Each droplet is therefore associated with two images: One cropped image of the droplet at full resolution (e.g. the images of Fig. 1c) and one low-resolution image of the complete chip. These images are useful in different stages of data curation.

To account for the relation between the different images of each droplet the data are structured using the relational database SQlite. The nested data structure is shown in Fig. 2, which shows how each droplet can be identified within each microfluidic chip, which itself is part of a multiple chip dataset. The droplet, chip, and dataset id are the same for days one and two. Some of the the metadata are common for each dataset, e.g. the type of antibiotic, while other metadata are specific to the chip level, e.g. the antibiotic concentration. Since the droplet id is also associated with its x-y position within the chip, it is possible to relate each cropped image with the mask within the full-chip image.

**Figure 2:**
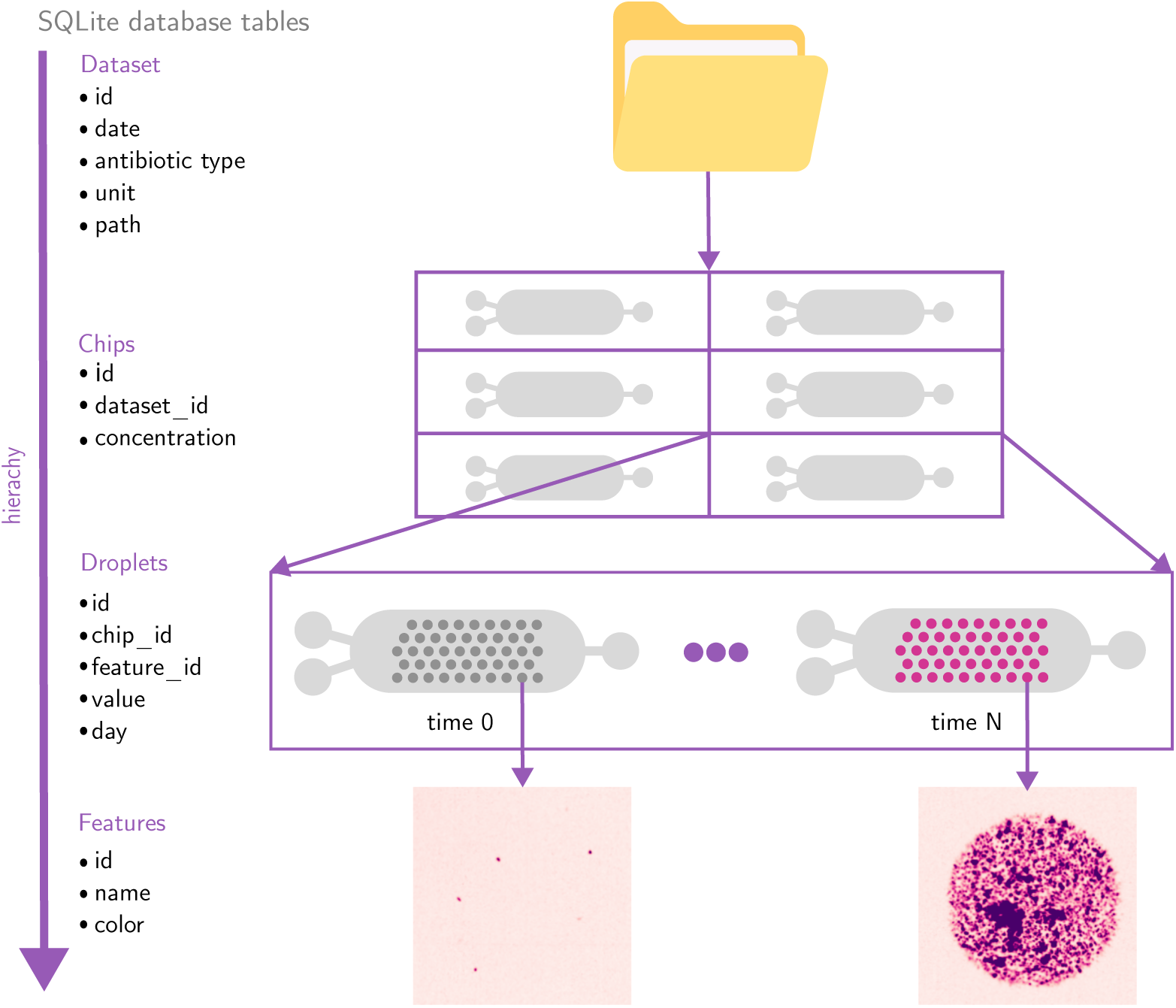
Hierarchical SQlite database structure. The database allows us to relate each droplet image to a given chip on a given time-point. The database can be updated dynamically through a web-based application.

Structuring the data in this way makes it possible to add information to each droplet in an interactive manner. This data curation is implemented through a web-based platform that fetches the images and meta-data from a server and allows the user to add labels to them directly (see Fig. 3). Two levels of interactions are implemented: On the first level the web application displays the low resolution image of the entire chip in the browser (bright-field and fluorescence), allowing the user to zoom and pan through the image interactively (Fig. 3a and supplementary movie S1). Every droplet in this image is highlighted with a grey circle. The user can then select a feature from the menu and assign it to a chosen droplet by clicking on it. Clicking a second time on the droplet removes the feature assignment. The web interface also offers other functionalities, including links to other concentrations from the same experiment, the ability to toggle between day 1 and day 2, and an option to display full-resolution images of the droplet for both days simultaneously when hovering with the cursor.

**Figure 3:**
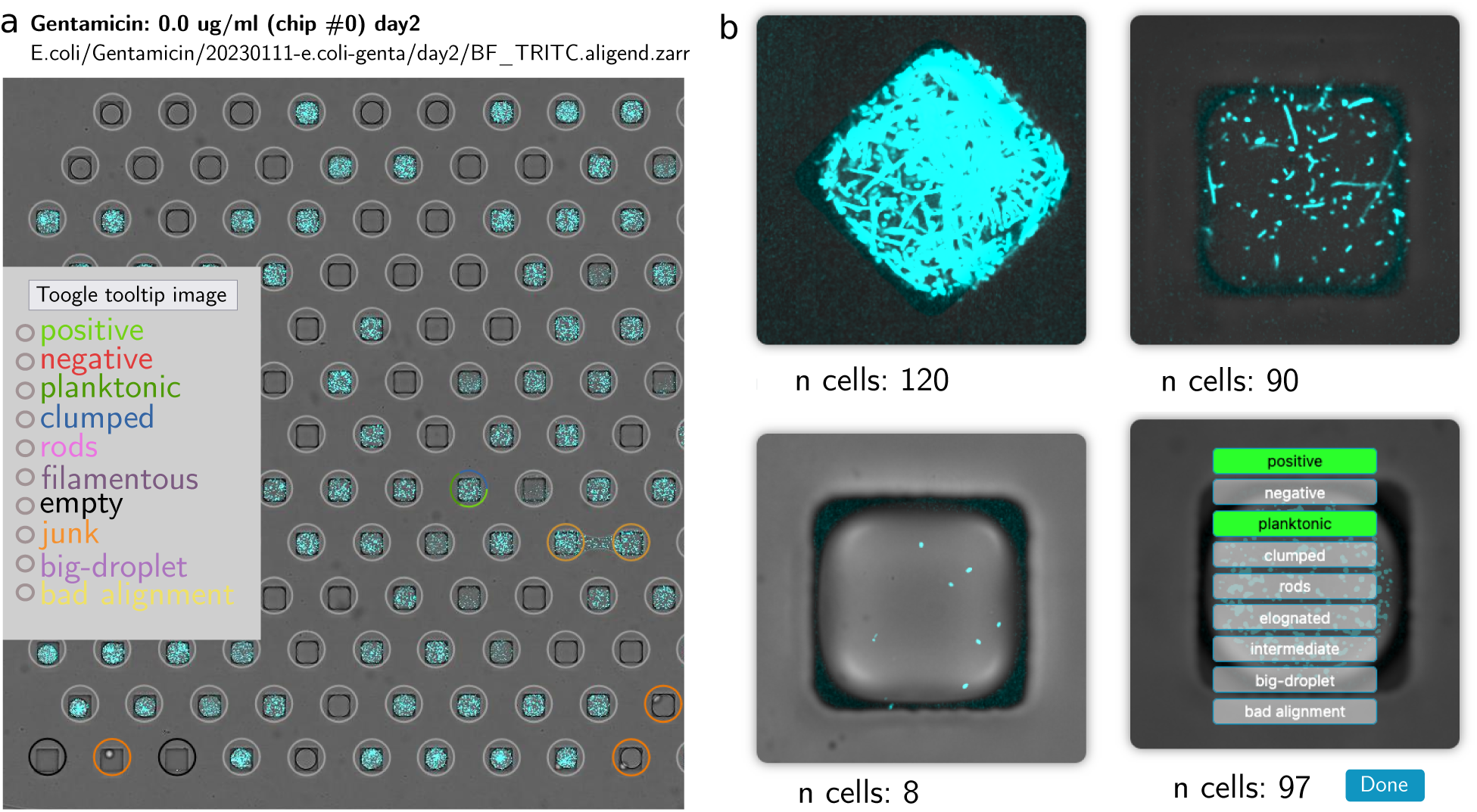
Interactive labeling of the droplets in context aware or context free modes. **(a)** Context aware interface, showing the whole chip at low resolution for labelling of bad droplets using bright-field. See supplementary movie S1 for interactions with the images. **(b)** Context free interface showing randomly selected droplets at full resolution. Used for morphology labelling. See supplementary movie S2 for interactions with the images. Note that the cell count is inaccurate for values *n* ≳ 100 (*9*).

This full chip view is context dependent and it is used for quality control and to exclude droplets from subsequent analysis. Situations that are excluded include positions that do not contain a droplet, drops that span several anchors (merged droplets), or fluorescent debris on the chips. In other cases droplets are excluded if they are too large (inconsistent breaking) or too small (due to evaporation). Finally, in some rare cases specific regions of anchors do not align with the predicted anchor position, resulting in bad image crops. This quality control step, which takes less than 5 minutes per experiment, allows us to ensure that the data that are later analyzed are of high quality.

The second level for droplet annotation is to identify the morphology of their contents in a context-free manner, i.e. without having the information of the antibiotic type, concentration or growth behavior of surrounding droplets. In this interface, high-resolution images of individual droplets are randomly selected from different experiments and displayed on the screen (see Fig. 3b and supplementary movie S2). The user can then add predefined labels to individual images, which are then used to generate a training dataset of colony morphologies. Previously labelled features are shown as activated buttons to prevent data duplication. Note that this manual annotation can be combined with automatic feature detection, such as cell number counts within each droplet. The database and interface are available as a docker image as described in data availability statement.

### Training and validation of the neural network

Input from the web-based interface was used to train a neural network to label the contents of each droplet. The curated dataset consisted of 1184 droplets, which were split 70:30 into a training and test set having a comparable label distribution between them. The distribution of morphologies are shown in Fig. 4a, both as a ratio to the total number of labelled drops and as a ratio of the total number of labels, since a single drop can have several labels. To perform the automated labelling task, pytorch’s convolutional neural network resnet34 with 34 layers was trained, using transfer learning: the network’s weights were initialized by default weights obtained originally by training on the ImageNet dataset (*30*). A weighted binary cross entropy with logit loss was used as an optimization function, see materials and methods (*31*).

**Figure 4:**
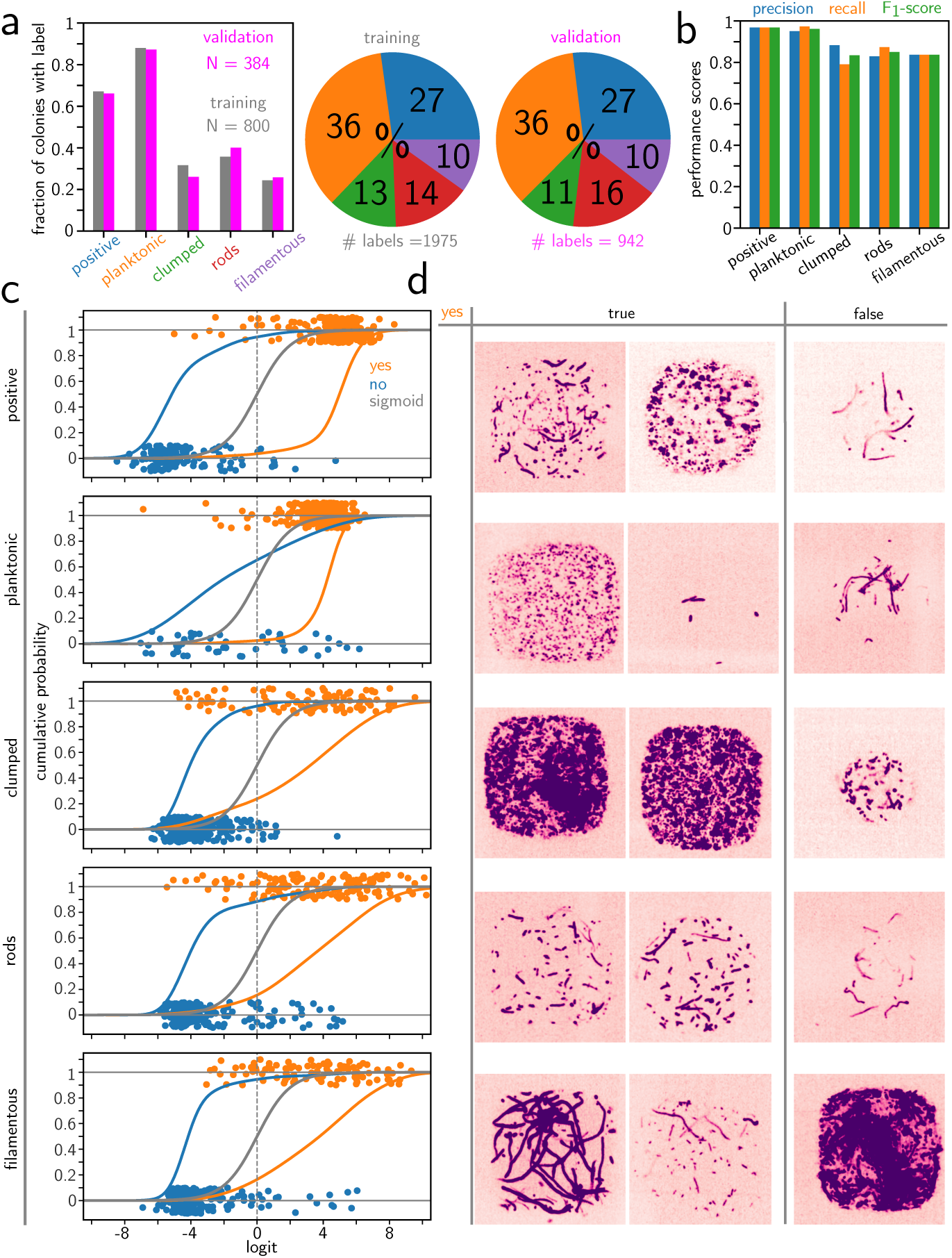
Machine learning allows automatic droplet labeling. **(a)** Distribution of morphologies for the training and validation datasets. Bar graph shows the fraction of droplets from total number of droplets *N*. The pie chart shows the fraction of labels, since labels are not mutually exclusive. **(c)** Summary statistics for precision, recall and *F*_1_-score for the network performance on the validation set. **(c)** Probability of the validation sets logits, grouped by the ground truth label (yes:orange,no:blue). The actual logit values are plotted as scatter with a jitter. The colored solid lines represent the corresponding cumulative probabilities. The gray line represents the sigmoid used to convert between logit and probability of the label. **(d)** Example images after classification. The two left images show examples of ”true positives” (yes, true), while the right-most image shows a ”false positive” (yes,false).

The training is evaluated by computing the average loss and subset accuracy after every epoch for the training and validation datasets. Both the average training and validation losses decrease and converge, indicating proper learning of our model (see Fig. S1). At the same time the subset accuracy of the training and validation sets increase. The training set achieves a near perfect score while the validation set’s subset accuracy saturates at around 65%. Since however the subset accuracy requires perfect prediction of every label on an image, which is very restrictive, precision, recall and *F*_1_-score are considered as suitable metrics for non exclusive multi label training (*32*). These values at the end of the learning are plotted in Fig. 4b, showing that the positive and planktonic archive scores are above 95% while the other labels reach values of at least 80%. In particular recall and precision are similar, indicating that we detect labels if they exist and also classify them correctly.

To better assess when droplets are correctly or incorrectly predicted we turn to the predicted logits of the validation set (see Fig. 4c). The logit is transformed into a label probability with the sigmoid function, such that a zero logit marks the point where the probability of a class label reaches 50%. In this figure the cumulative probability of the logits is plotted to provide better visualization. It shows that values with a ”no” label (blue data) cluster at negative values of the logits, indicating that the network identifies the absence of a morphology correctly. This result is less pronounced for the ”planktonic” label, for which the network had only few examples of truly non planktonic colonies to train on.

For images with a ”yes” value (orange data) the cumulative probability at zero logit corresponds to 1 minus the recall. However, the cumulative probability contains more information. For positive and planktonic labels the majority of the labels is concentrated at larger logits, indicating strong recognition by the neural network. This is also highlighted in their precisions and recalls that are larger than 95%. The other labels show less pronounced clustering but still have recalls of above 80%.

Some example images of true positive and false positive labels are shown in Fig. 4d, where we focus only on droplets containing a single label for clarity. We observe that images for which the network falsely predicts the existence of a label correspond to cases where the ground truth label is not obvious even for the human eye (see Fig. 4d). A different false positive classification is often seen for rods, where the human marks the short bent cells as filaments but the network predicts them to be rod-like. For the filamentous labels we observe false positive labels from colonies that show dense regions which we decided to mark as clumped.

### Bacterial morphology types for different antibiotic concentrations

The neural network was used to identify the contents of all non labelled droplets, which yielded labels on a total of 82,268 droplets over all antibiotics and concentrations, as shown in Fig. 5. We first verify that the fraction of positive droplets which decreases with increasing antibiotic concentration, as expected. Note that this decrease is regular and reproducible for most cases except for chloramphenicol, for which different experimental replicates produce widely varying results.

**Figure 5:**
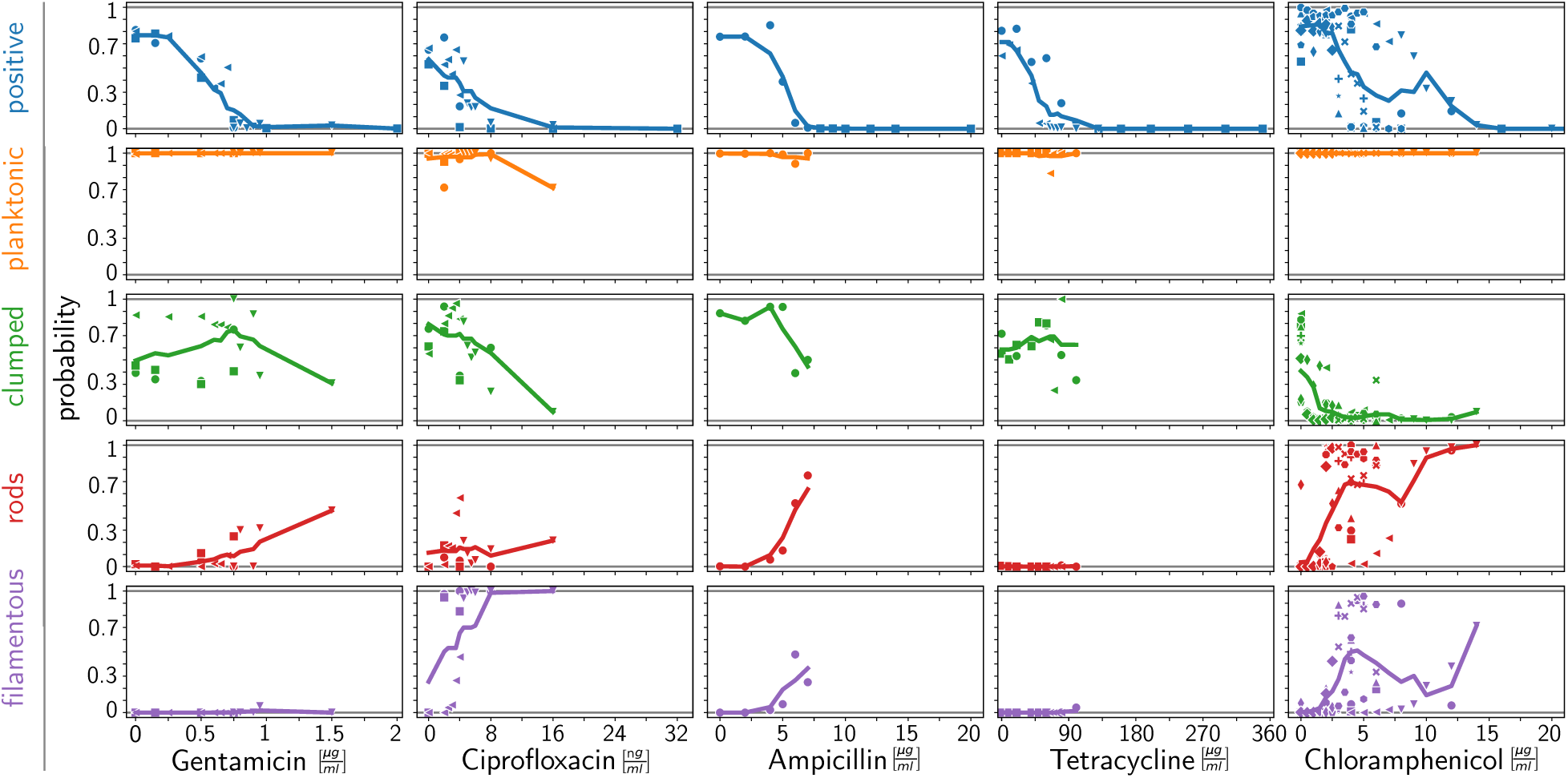
Morphological profiles of surviving cells by antibiotic type. **(a)** Fraction of of positive droplets, and fraction of other morphologies conditioned on being a positive droplet as a function of concentration and type of antibiotic. Different symbols correspond to different biological replicates. The solid lines show the moving average with window sizes 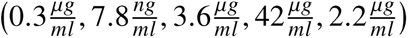 respectively. The window sizes correspond to the weighted concentration averages, with the number of bacterial cultures as weights. Only conditions where the number of positive droplets is larger than two are considered.

Now that the positive droplets are individually identified, the rest of the morphology quantifications in Fig. 5 are shown conditionally for positive droplets (for the unconditioned profiles see SI. Fig. S2). The data show that the planktonic phenotype is detected in nearly all positive droplets, for all antibiotics, except for a small decrease at high concentrations of ciprofloxacin. Moreover clumping is also frequently observed to coexist with the planktonic case, including in the absence of antibiotics. This clumping behavior is consistent with well-documented observations of adhesion between individual E. coli cells, which helps determine colony morphology and growth (*33*) while also impacting the biological function of the cells (*34*). A notable exception to the clumping phenotype is the case of chloramphenicol, where the clumping morphology nearly disappears even for very low concentrations of antibiotic, suggesting that the drug inhibits the adhesion molecules on the cell surface. Beyond these generally present morphologies, rods and filaments appear for the different antibiotics to varying degrees, except for tetracycline which does not form elongated shapes. Gentamicin only produces short and straight rods, while the ciprofloxacin morphology is dominated by long bending filaments. Ampicillin and chloramphenicol display both rods and filaments.

Compared with previously reported morphology changes under antibiotic stress, the breadth of the measurements covered in this study provide a more standardized classification. For instance it has been reported that ampicillin and ciprofloxacin lead to filamentation in E. coli (*23, 35*) and chloramphenicol leads a moderate increase in cell length and cross-section (*36*). From the current measurements it emerges that ampicillin-associated filaments are short rods, while those under ciprofloxacin are long and curved filaments. Similarly, previously reported elongation under gentamicin (*16*) corresponds to rods in our classification. In contrast with the cases above, rods and filaments often coexist in the case of chloramphenicol.

It is interesting to contrast the results for tetracycline and chloramphenicol. Even though both antibiotics target the protein synthesis, by binding to the 30S and 50S ribosomal subunit respectively, the associated phenotypes are very different. Tetracycline does not display a modification of phenotype compared with the case for no antibiotic, while chloramphenicol loses its clumping ability and displays rods and filaments even for low concentrations. These differences in morphological signature indicate that chloramphenicol leads to a strong modification of the cellular functions that is not observed for tetracycline.

### Multidimensional digital signatures

The results for different antibiotics can be compared together on a common scale by replacing the antibiotic concentration with its potency in each microfluidic chip, as shown in Fig. 6a. This potency is measured by counting of the number of negative vs. positive droplets within each chip and computing the probability of a droplet to be negative, thus combining the different digital measurements of survival and morphology. In addition to providing a scale to compare different antibiotics together, this normalization provides an internal scale for each chip and thus reduces the reproducibility issues that are often encountered with microbiology experiments.

**Figure 6:**
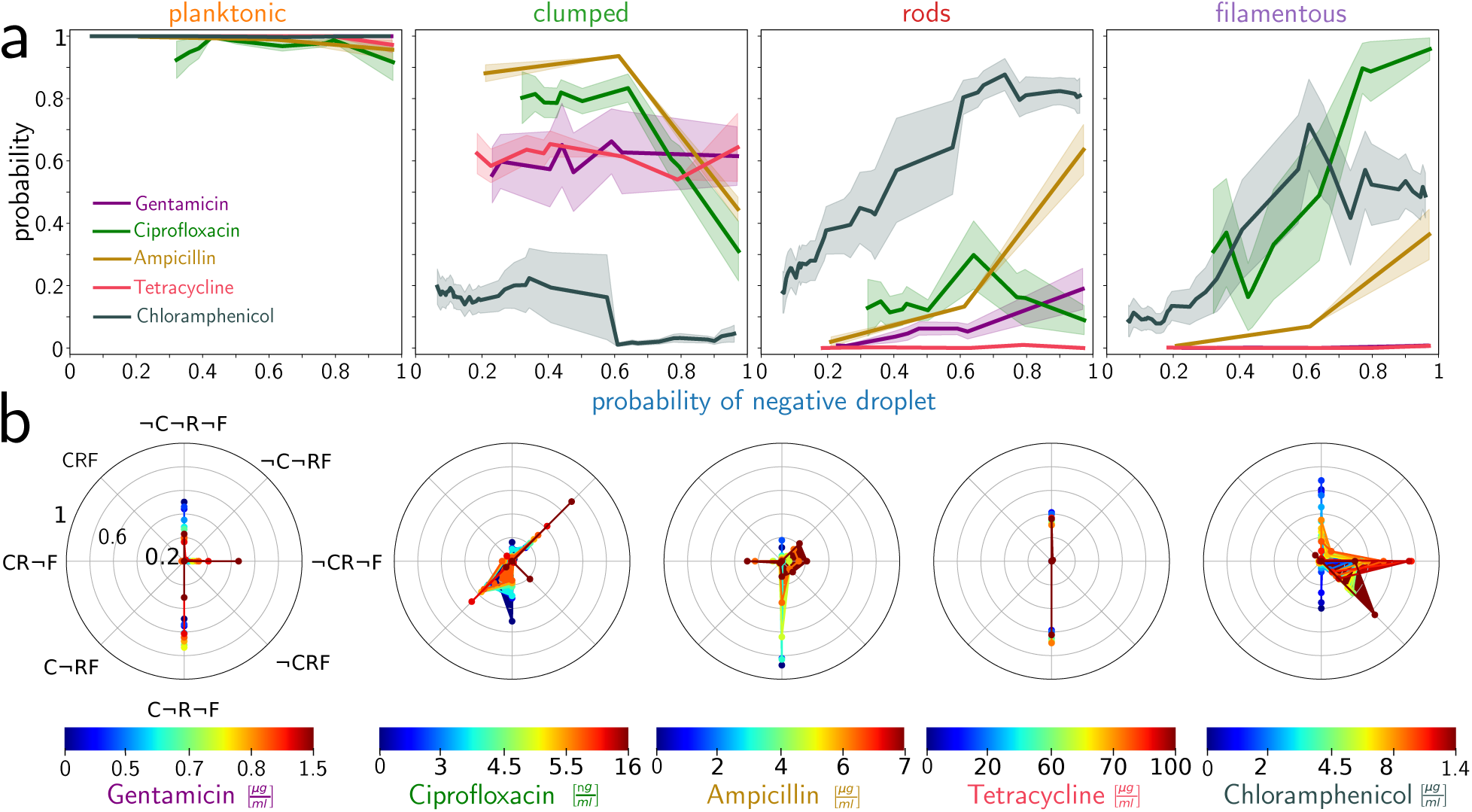
Multidimensional morphological profiles provide unique signatures for each antibiotic condition. **(a)** Evolution of the different morphologies as a function of the probability of droplets to be negative. The lines correspond to running averages with a window size of 0.25, while the shaded area corresponds to the SEM. **(b)** Co-occurrence probability of morphologies conditioned on positive colony formation for each antibiotic. C,R and F represent the existence of clumped, rod-like and filamentous cells respectively. The ¬ symbol before the letter marks the absence of the given phenotype. The colour scale is a non-uniform scale, and chosen to help with visibility.

The graphs on Fig. 6a address the key question of knowing which morphologies are selected by the surviving cells as the conditions for growing into a colony become more difficult. It emerges that the rod and filament morphologies appear at different killing probabilities and grow at different rates for different antibiotics, indicating that the stress that leads to these morphologies is indirectly linked with the killing capacity of the drug. Moreover, the disappearance of the clumped phenotype occurs quickly for chloramphenicol but more slowly for ampicillin and ciprofloxacin, but stays constant for tetracycline and gentamicin.

Finally the encapsulation within droplets allows a multi-dimensional signature to be computed that account for the co-existence of the different morphologies within a single droplet (Fig. 6b). Since the labels are not mutually exclusive, each droplet can be characterized by one, two, or more features. It is therefore possible to plot the probability for each drop to contain the combination of morphologies as a function of antibiotic type and concentration. The resulting graphs, shown in Fig. 6b, show that each antibiotic presents a distinct signature as a function of concentration. Compared with previous measurements that only identify features independently of each other, the combination of features allow us to identify which morphologies occur together and which are mutually exclusive, for each condition. These results are particularly interesting a high antibiotic concentrations, for which only a small number of cells are expected to survive. In this case the multiple morphologies can be traced to the progeny of a single mother cell, showing that the successive divisions can lead to a population with a combination of different morphologies.

## Discussion

The measurements shown here were enabled by the simultaneous development of the microfluidic, microscopy, and software aspects. Now that these elements have been individually validated and integrated together, the workflow can be scaled up to millions of droplet images, including for instance for time-lapse imaging. The hardware and software tools are provided on open source platforms and can now be used to address a variety of microbiology questions on a wide range of microorganisms, e.g. different bacterial or yeast strains.

By observing the ability to form colonies with the resulting morphologies in each droplet we can directly link the diversity of morphologies to survival. This is of particular interest at high antibiotic concentrations where only few cells, which are of particular interest, survive. The use of the potential for colony formation as internal scale allows us to compare different antibiotic conditions on a unified scale and judge the diversity changes.

Compared with previous works that characterize the morphology of bacterial colonies (*37, 38*) or of individual cells (*36, 39, 40*), the results obtained here characterize a combination of scales, including individual cell shape and their interactions in the form of clumping and ability to divide. This *”collective scale”* characterization captures the impact of the antibiotic on the intra-cellular processes, which determines their shape, and on their interactions, which determines how they organize together. Just as characterizations at this collective scale have already provided useful insights into E. coli communication (*41*) or migration (*42*), they are likely to play a role in explaining aspects of the resistance to antibiotics, which is mediated by local population dynamics (*43*).

## Supporting information

Context aware labeling within microfluidic chip

Context-free labeling of microfluidic droplets

## Acknowledgments

The authors acknowledge the support of the Biomaterials and Microfluidics platform at Institut Pasteur. The bacteria were kindly provided by Julia Bos from Institut Pasteur.

## Author contributions

L.Le Q., A.A. and E.M. performed experiments. A.A. developed the web platform and E.M. and L.Le Q. performed labelling. E.M. performed machine learning, subsequent data analysis and wrote the original draft. C.N.B. supervised the research and wrote the original draft. E.M. A.A. and C.N.B. discussed the results, reviewed and edited the manuscript.

## Data and materials availability

All data needed to evaluate the conclusions in the paper are present in the paper and/or the Supplementary Materials. The imaging data and sql database is available at on Zenodo with the identifier 10.5281/zenodo.17193389 (*44*). The processed data are composed of csv tables and python jupyter notebooks that allow the production of all figures in the paper. These are all available at Github (*45*). The github repository also contains instructions on how to access and run the web platform as a docker container, for the inspection of the images and training annotations.

## Supplementary Materials for

### Materials and Methods

#### Cell culture and loading preparation

##### Strain

In all experiments the E. coli W3110 strain labelled with 405 red fluorescent protein (lacYZ:mRFP-1) was used (*29*).

##### Cell culture

From the -80°C stock, the cells were streaked on LB agar plates and incubated overnight at 37°C. The next day one isolated colony is inoculated in lysogeny broth (LB) and, to induce the expression of RFP, IPTG is added at 0.05 mM. The bacterial suspension is then incubated overnight at 37°C while shaking.

##### Cell dilution

To prepare the cell solution for loading, we measured the optical density (OD) at 600 nm and dilute the sample into fresh cell culture medium (LB + IPTG) to a target OD of 10^−3^, resulting in approximately 0-10 cells per droplet.

##### Antibiotic Solution

Gentamicin (Sigma-Aldrich) was solubilized in MilliQ water at 50 mg/mL. The stock was then diluted with MilliQ water to 5 mg/mL.

Ciprofloxacin (Sigma-Aldrich) was solubilized in MilliQ 0.1 N HCl (Sigma-Aldrich) at 25 mg/mL. The stock was then diluted with MilliQ water to 1 *μ*g/ml.

Chloramphenicol (Sigma-Aldrich) was solubilized in ethanol at 10 mg/mL. The stock was then diluted with MilliQ water to 5 mg/mL.

Tetracycline (Sigma-Alrich) was solubilized in MilliQ water at 50 mg/mL. The stock was then diluted to 1 mg/mL.

### Loading

To load the cell-antibiotic-solution we use a NEMESYS syringe pump and the protocol described in (*9*). The chip is first filled with continuous oil phase consisting of 3M FC40 fluorinated oil and RAN fluoSurf surfactant at 1%. After purging all air bubbles, the chip is loaded with the combined bacterial-antibiotic solution and again filled with the continuous oil phase to induce droplet formation. The loaded chip is then put on ice and in the dark to limit cell growth and photobleaching. Once all chips are loaded they are transferred to the microscope.

### Microscopy and Image Analysis

We use the same Microscopy-Image analysis pipeline as detailed in (*9*). Microscopy images are acquired using spinning disk confocal microscope (Nikon Ti2 + Yokogawa) with a 20x 0.7 NA air objective lens (Nikon Inc.) and with 215×2 pixels binning (set directly in camera properties (Hammamatsu Orca 4)). Images of the complete chip are obtained by stitching individual images with a 5% overlap. We acquire first a bright-field image of the complete chip, and the RFP signal is imaged in confocal mode, using 3D stacks with 5 *μ*m steps for a total penetration depth of about 120 *μ*m. The imaging is followed by maximum projection, image registration and cell counting.

### Interacting with images in the browser

To interact with the images in the browser we need to render the images which are stored in multiscale zarr arrays. To load a particular chip the backend uses a universally unique identifier (uuid) to retrieve the image path from a SQLite database. For fast rendering, the brightfield and fluorescent images are stored with 815×8 binning, which the backend converts to a 24 bit RGB image that is encoded as a base64 string, that can be part of a JavaScript Object Notation (json) object. The backend also reads an associated csv which contains the droplet coordinates and retrieves the existing labels and cell counts from the database, which are all sent to the browser as an json. The canvas of the browser renders the image on the screen and uses the droplet coordinates to draw gray circles. The circles shown around droplets act as interactive regions, where the canvas detects if a cursor click occurs. As described in the main text this feature is used to assign labels from the left menu, which then change the colour of the drawn circle according to the selected features. At the same time the browser saves the new label to the database. The backend checks if this label already exists for this chip and either adds a new record or deletes the existing one. The user can download a csv file containing the labels.

The interface also allows to activate a hover functionality, where the canvas will show a full-resolution crop of the droplet for day 1 and day 2 simultaneously, while the cursor hovors over the droplet. These images are retrieved from a separate zarr array containing only the cropped images of the individual droplets in original resolution.

Additionally, we implemented a context free labelling interface where a few droplets crops of size 300 x 300 pixels are shown without metadata about the experiment (Fig.3b). In fact the droplets shown are randomly selected from the database and can come from many antibiotics and concentrations. These droplets are aligned with the labelling buttons and are calling the same API endpoints as the context aware interface to save labeles into the database.

### Machine learning

We use pytorch’s resnet34, a convolutional neural network with 34 layers and apply transfer learning were we initialised the network by default weights that have originally been obtained by training on the ImageNet dataset. To comply with the requirements of this dataset we transform all our images to RGB images, resizing it to 24415×244 pixels and normalising the images to means values (0.485, 0.456, 0.406) and standard deviations (0.229, 0.224, 0.225). Our loss optimisation function, is binary cross entropy with logit loss, were we load batch sizes of 32 images, dropping incomplete batches, and applying randomly horizontal and vertical flips before feeding the images into the network. The loss minimisation is performed by the adam solver with initial learning rate of 10^−5^, and a learning scheduler with period of 50 epochs and factor 0.1 for learning rate decay. We optimise over 100 epochs and compute the average loss and subset accuracy (percentage of samples that have all their labels classified correctly) during training for both the training and validation datasets. At the end of training we asses the recall 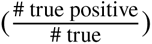 precision 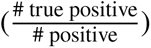, and their harmonic mean the *F*_1_-score of the validation dataset per label.

### Supplementary Text

**Figure S1:**
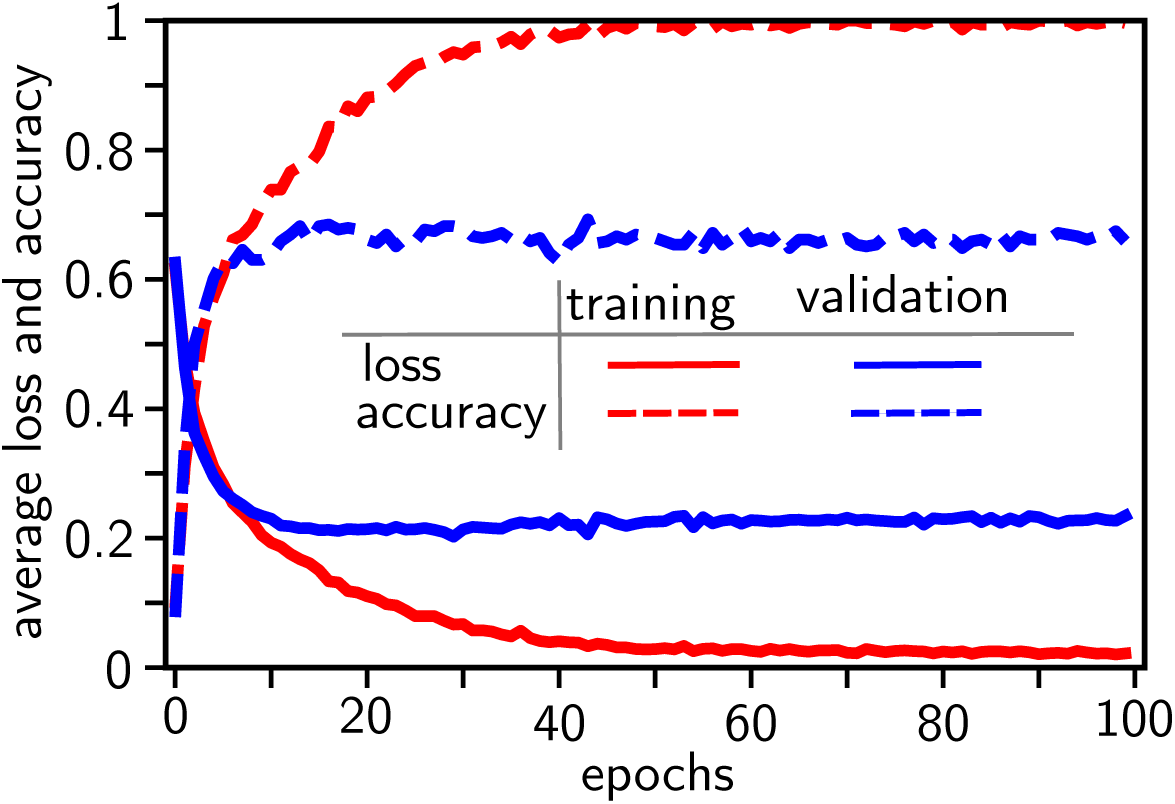
Supervising the machine learning. Loss and subset accuracy of test and validation dataset.

**Figure S2:**
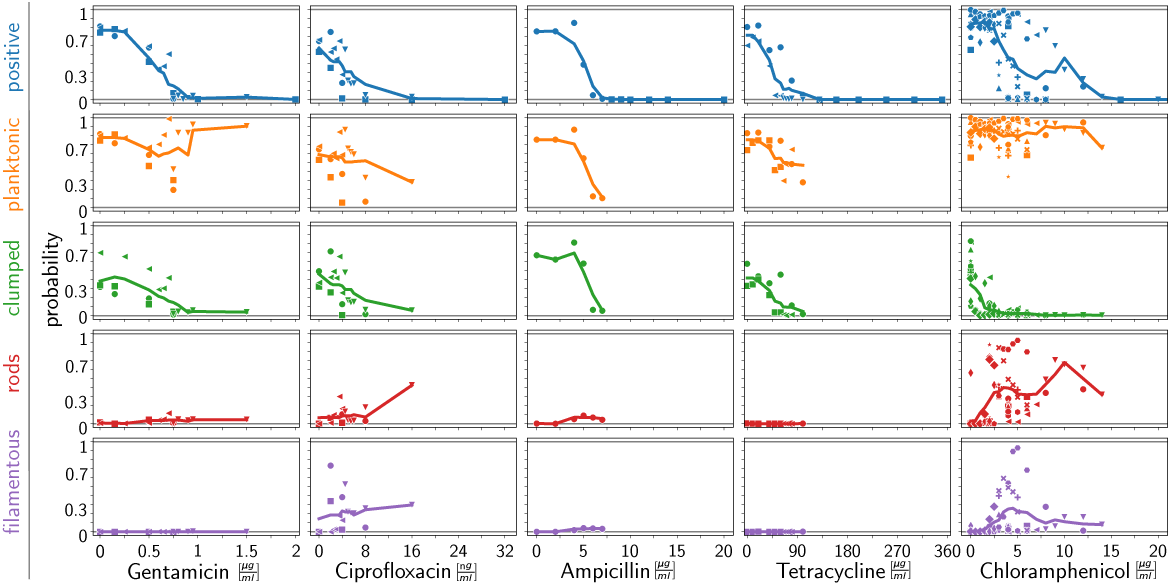
Unconditioned morphological survival profiles. Fraction of morphologies as a function of concentration and type of antibiotic. Different symbols correspond to different bacterial cultures, and the solid line shows the moving average with window sizes 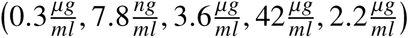 respectively. The window sizes corresponds to the weighted concentration averages, with the number of bacterial cultures as weights. Only conditions where the number of positive droplets is larger than two are considered.

**Caption for Movie S1. Context aware labeling.** Illustration of the context aware labeling platform, that allows to switch between datasets, show full resolution crops of individual droplets and the ability to label them.

**Caption for Movie S2. Context free labeling.** Illustration of the context free labeling platform that randomly displays droplets from the database and allows labeling. Previous set labels are shown to avoid duplication.

